# scRNAseq of healthy and irradiated mouse parotid glands highlights crosstalk between immune and secretory cells during chronic injury

**DOI:** 10.1101/2022.11.26.517939

**Authors:** Brenna A. Rheinheimer, Mary C. Pasquale, NIDCD/NIDCR Genomics and Computational Biology Core, Kirsten H. Limesand, Matthew P. Hoffman, Alejandro M Chibly

## Abstract

Translational frameworks to understand the chronic loss of salivary dysfunction that follows after clinical irradiation, and the development of regenerative therapies remain an unmet clinical need. Understanding the transcriptional landscape long after irradiation treatment that results in chronic salivary hypofunction will help identify injury mechanisms and develop regenerative therapies to address this need. Advances in single cell (sc)RNAseq have made it possible to identify previously uncharacterized cell types within tissues and to uncover gene regulatory networks that mediate cell-cell communication and drive specific cell states. scRNAseq studies have been performed for virtually all major tissues including salivary glands; however, there are currently no scRNAseq studies evaluating the long-term chronic effects of irradiation on salivary glands. Here, we present scRNAseq from control and irradiated murine parotid glands collected 10 months post-irradiation. We identify a population of epithelial cells in the gland defined by expression of *Etv1*, which may be an acinar cell precursor. These Etv1+ cells also express *Ntrk2* and *Erbb3* and thus may respond to myoepithelial cell-derived growth factor ligands. Furthermore, our data suggests that CD4+CD8+ T-cells and secretory cells are the most transcriptionally affected during chronic injury with radiation, suggesting active immune involvement during chronic injury post-irradiation. Thus, our study provides a resource to understand the transcriptional landscape in a chronic post-irradiation microenvironment and identifies cell-specific pathways that may be targeted to repair chronic damage.

**Highlights:**

- We generated a scRNAseq dataset of chronic irradiation injury in parotid glands
- A newly identified *Etv1*+ epithelial population may be acinar precursors
- Ntrk2 and Erbb3 are highly specific Etv1+ cell receptors that may mediate cell-cell communication with myoepithelial cells
- CD8+ T-cells and secretory acinar cells have the greatest transcriptional changes post-IR

## Introduction

Of the three major pairs of salivary glands (SGs): the parotid (PG), submandibular (SMG), and sublingual (SLG), the human PG is the largest and produces the largest volume of saliva, particularly in response to gustatory simulation. In mice, the PG is smaller than the SMG but also contributes to the majority of stimulated saliva ^1^. In addition, the PG is the most sensitive to irradiation (IR) damage, a therapeutic treatment for head and neck cancer that often results in permanent salivary hypofunction. In terms of understanding salivary gland biology, most studies have focused on the SMG both in the context of development and response to injury; however, each gland has unique functions and transcriptional profile ^2^. Here we set out to investigate the effects of irradiation damage to PGs in mice using single cell (sc)RNAseq.

The PG is primarily comprised of serous acinar cells which produce large volumes of watery serous saliva that is transported through the ductal system into the oral cavity to aid in digestion and protection of mucosal surfaces. Despite advances in radiotherapy, it is estimated that ∼40% of head and neck cancer patients suffer from the chronic consequences of salivary gland damage months to years after the completion of radiotherapy, even with newer modalities such as intensity-modulated radiation treatment (IMRT) that reduces exposure to non-tumor tissues ^3–7^. Animal studies show that the acute effects of radiotherapy in the PG occur in the days and weeks following initial treatment and are likely a result of high levels of acinar cell death ^8^, DNA damage ^9^, dysregulated calcium signaling and ROS generation ^10^, inflammatory responses ^11^, and alterations to the nerves and vasculature, whereas the chronic effects arise months to years after initial treatment (Reviewed in ^12^). Chronic loss of function is often attributed to fibrosis and the inability of acinar regeneration to occur, and preclinical studies suggest that persistent acinar cell proliferation, vascular damage, and parenchymal cell loss may be contributing factors ^13–16^. In a similar manner, patients with Sjogren’s syndrome, an autoimmune disease that damages the acinar cells of salivary and lacrimal glands, life-long consequences include dental caries, reduced taste and smell, malnutrition, mucositis, and increased risk for oral infections leading to a significant decrease in quality of life ^17^. Therefore, translational frameworks to understand chronic glandular dysfunction following IR therapy along with the development of regenerative therapies remains an unmet need.

The development of scRNAseq has made it possible to identify previously uncharacterized cell types within a tissue and to uncover and gene regulatory networks and mechanisms regulating cell-cell communication and specific cell states ^18–21^. To date, there have been scRNAseq studies performed for virtually all major tissues, including atlas-level scRNAseq datasets such as the Tabula Muris ^22^ or the Tabula Sapiens ^23^ which integrate data from multiple organs in mouse and human, respectively. There are also numerous scRNAseq studies on disease-specific models, which are important to understand the cellular mechanisms involved that could be targeted for repair or regeneration. In SGs, scRNAseq studies have focused on understanding homeostasis and development ^24–28^, as well as particular disease states, such cancer ^29^, Sjogren syndrome ^30, 31^, and COVID-19 infection ^25^.

In this study, we use scRNAseq to characterize the adult mouse PG and compare the transcriptional landscape 10 months after IR damage to explore chronic dysfunction post-irradiation. Due to the complex heterogeneity of the SGs, distinguishing cell-type compositional differences and their specific and direct contribution to the loss of saliva following radiation therapy is complex, and single-cell transcriptomics will begin to resolve this issue.

This dataset allows for discovery and exploratory research into the mechanisms and cellular processes driving PG dysfunction post-IR in a model of fractionated IR with limited acinar cell loss. Our work has been validated by immunofluorescence staining to confirm the presence of selected markers in specific cell populations, confirming the potential to reveal meaningful biological insights. It is noteworthy that scRNAseq of in vivo models of chronic IR injury has only been performed in liver ^32^, lung ^33^, and skin ^34^, and data is only publicly available for lung and skin. Thus, our study will also be an essential resource to better understand cell-specific responses to IR in general.

## Results

### Generation of a single-cell resource of healthy and irradiated mouse parotid gland

Using the 10X Genomics platform, we generated 2 individual scRNAseq libraries of healthy and IR mouse PG collected 10-months post-irradiation (Figure 1A). Mice received 6 Gy IR/day to the head and neck region on five consecutive days, for a total dose of 30 Gy. This mouse model of IR damage to SGs results in chronic loss of saliva with partial loss of epithelial cells ^35^. Control and IR PG samples were bioinformatically integrated with SEURAT v4 and clustered following SEURAT’s standard workflow ^36, 37^. The optimal resolution for clustering was determined using clustree package ^38^ and the resulting 17 cell clusters were annotated based on their gene expression profile (Figure 1B, S1A-B) and a previously generated atlas of SMG development which provided cell type specific markers ^26^. Stromal and myoepithelial cells (MECs) clustered together with endothelial cells likely due to the low number of cells recovered for these populations. Thus, they were manually annotated based on expression of a combination of stromal (*Col1a12* and *Vim*) and myoepithelial (*Krt14* and *Acta2*) markers which were highly specific (Figure S1C-D). We did not identify discrete clusters of basal duct cells (*Krt14+Krt5+*) or peripheral nerves presumably due to limitations in the dissociation technique, which has been previously reported for adult SG tissue dissociation ^26^.

**Figure 1.**
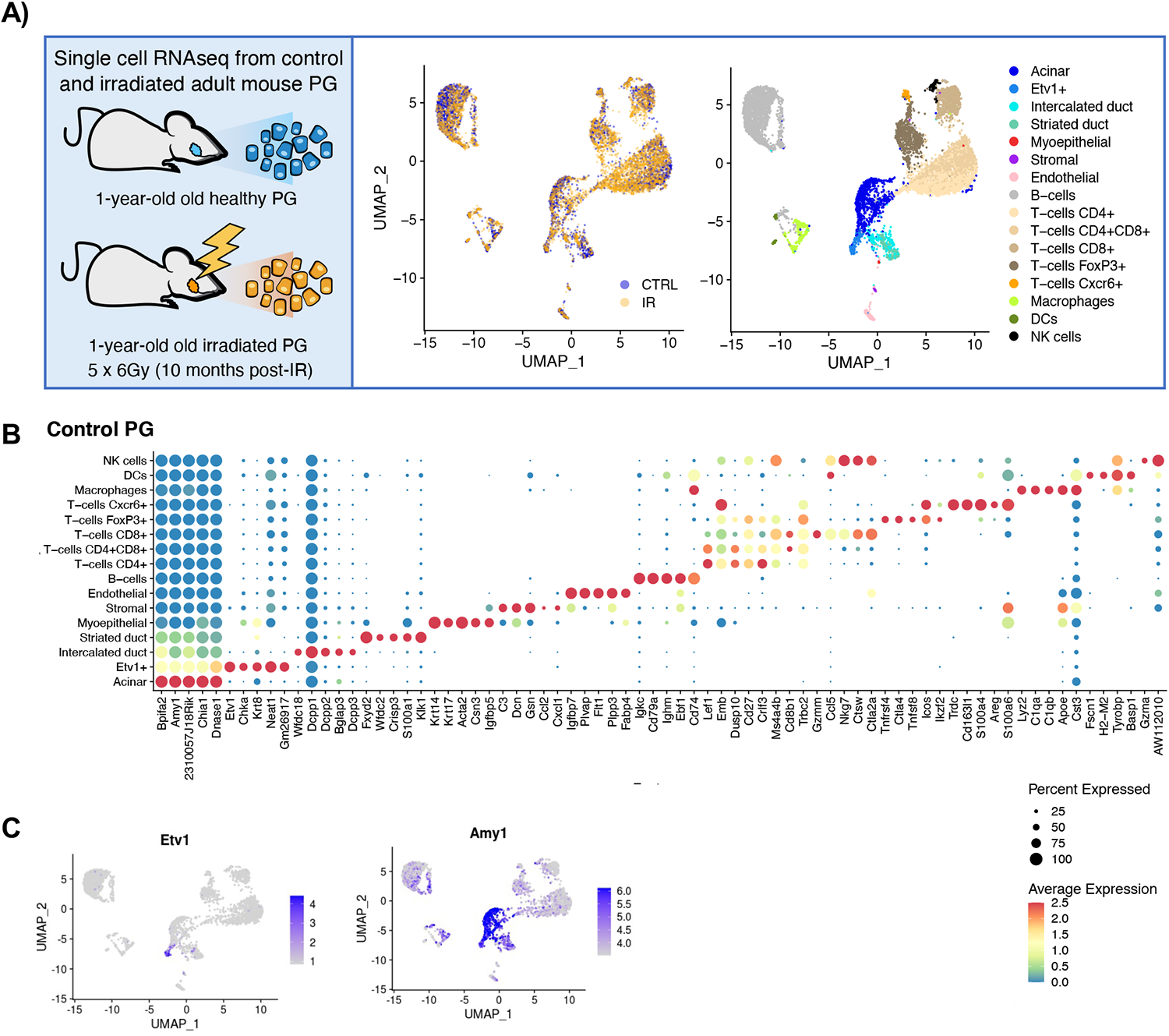
scRNAseq analysis of control and irradiated PG. A) Single cell suspensions from 1-year-old control and irradiated PG from 2 C3H female mice were used to build scRNAseq libraries. Representative UMAP plots are colored by treatment group or cell type. Clusters were annotated based on the expression of known markers. B) Balloon Plot with top 5 DEGs per cluster sorted by fold change. Statistical analysis performed using SEURAT package in R. Color is relative to scaled gene expression and size of the dot represents the percentage of cells expressing the gene. C) Representative UMAP plots showing expression of Etv1 and Amy1

The identified populations included acinar cells (*Amy1+*), intercalated duct (*Dcpp1-3+*), striated duct (*Fxyd2+, Klk1+*), MECs (*Acta2+Krt14+*), stromal (*Col1a1+Vim+*), endothelial (*Pecam1+*), and 9 distinct immune populations including B-cells (*Cd79a+* and Immunoglobulin genes), five subtypes of T-cells (*CD4+; CD8+; CD4+CD8+; FoxP3+; Cxcr6+*), macrophages (*Adgre1+*), dendritic cells (*S100a8/9+*), and natural killer cells (*Gzma+Nkg7+*). We also identified a previously uncharacterized epithelial population defined by high expression of *Etv1* and *Krt8* and moderate expression of *Amy1* (Figure 1B-C, S1B).

### Etv1 expression delineates a secretory subpopulation in acinar and duct compartments

The two most striking observations from our initial clustering analysis are the identification of an *Etv1*+ epithelial population and the prominence of multiple resident immune cell types after IR. *Etv1* has been described as one of the top transcription factors expressed in the salivary glands ^39^. To characterize this *Etv1+* cluster, and to generate gene expression profiles of individual cell populations in healthy adult parotid glands, we performed differential expression analysis with SEURAT in the annotated control sample (Figure 1C). Genes enriched in a given cluster are herein referred to as cell-defining genes and were sometimes expressed elsewhere at lower levels. The complete gene list is included in Supplementary File 1.

The expression of Amy1 in *Etv1+* cells suggested an acinar-like phenotype. When comparing the gene expression profile of major epithelial populations, 38% of acinar-defining genes (30 of 79) were enriched in *Etv1*+ cells (Figure 2A-B). Both cell types expressed serous secretory markers such as amylase (*Amy1*), parotid secretory protein (*Bpifa2*), prolactin induced protein (*Pip*), and carbonic anhydrase 6 (*Car6*), but their expression was significantly higher in acinar cells, while *Etv1+* cells had higher expression of *Krt8, Krt18,* and *Phlda1* (Figure 2C). When compared to duct populations, *Etv1+* cells expressed 38% (19 genes) of intercalated duct (ID)-defining genes (Figure S2A) and only 9.3% of striated duct (SD)-defining genes (Figure 2B, S2B), suggesting that *Etv1*+ cells are transcriptionally similar to both acinar and ID populations. Accordingly, Etv1 protein was detected by immunofluorescence in a subset of duct and acinar cells. Duct cells showed strong nuclear and cytoplasmic Etv1+ signal while it was predominantly nuclear in NKCC1+ acinar cells (Figure 2D).

**Figure 2.**
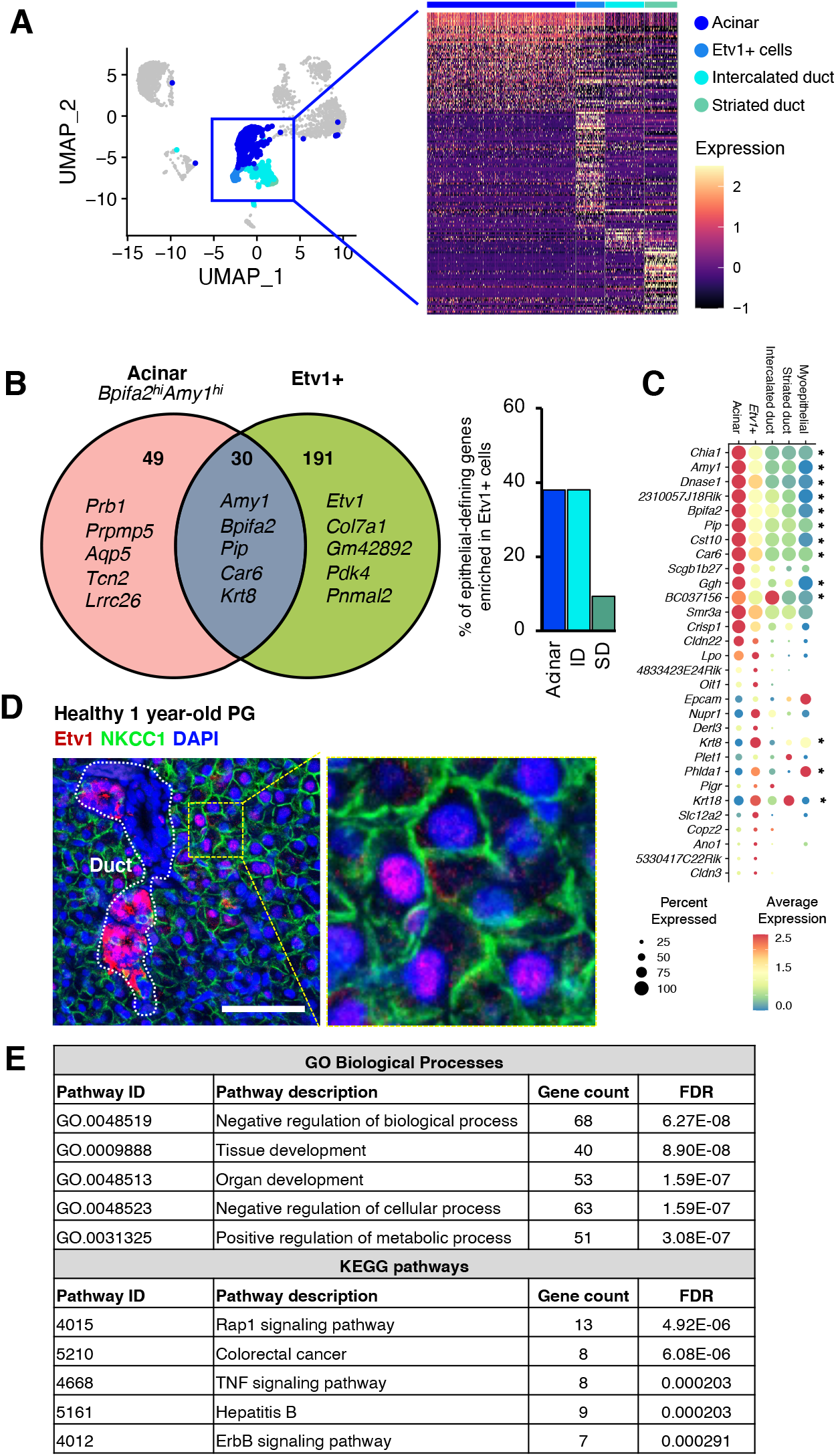
Characterization of acinar and Etv1+ cells. A) UMAP plot highlighting acinar, Etv1+, and duct populations with a representative heatmap of their gene expression profiles. B) Venn diagram of cell-defining genes in acinar and Etv1+ clusters showing the number of unique and overlapping cell-defining genes. Representative genes from each group are shown. The bar graph shows the percentage of overlap between cell-defining genes in acinar and duct populations with Etv1+ cells. C) Balloon plot showing expression of the 30 genes overlapping between acinar and Etv1+ cells. Genes marked with an asterisk are differentially expressed between Etv1+ and acinar cells (p<0.05, Wilcoxon rank sum test (SEURAT)). Color is relative to scaled gene expression and size of the dot represents the percentage of cells within a cluster expressing the gene. D) Immunofluorescence staining of PG from 1 year-old C3H mice stained for Etv1 (Red), NKCC1 (green) and DAPI (blue). The area delineated by the yellow dotted line is magnified to the right for visualization. Scale bar = 50um. E) Results from STITCH analysis showing top biological processes and KEGG pathways associated with defining-genes from Etv1+ cells.

Next, we performed functional analysis of all acinar and *Etv1*+ cell-defining genes using STITCH (search tool for interactions of chemicals, http://stitch.embl.de/), which integrates information about interactions from metabolic and KEGG pathways, crystal structures, binding experiments, and drug-target relationships. ^40^. As expected, KEGG pathway analysis on acinar genes showed salivary secretion as one of the top pathways (Figure S2D). In contrast, in *Etv1*+ cells the top functions and pathways were associated with organ development and activation of Rap1, TNF, and ErbB signaling pathways (Figure 2E, S2C), suggesting that the *Etv1+*population has distinct functions despite their transcriptional similarities to acinar cells.

### Acinar and Etv1+ cells communicate with MECs and stromal cells via Erbb3 and Ntrk2

Given that cellular functions are often initiated by ligand-receptor interactions that trigger signaling cascades, we next used two bioinformatic approaches to predict putative cell-cell interactions: First, we used differential expression analysis for each cluster and cross-referenced the resulting cell-defining genes with a previously published database of curated ligand-receptor pairs ^41^. For reproducibility of this approach, we used R-scripted code which is available as supplementary material. In this database, a ligand is defined as any molecule that interacts with known receptors and thus intracellular components such as *Hras* are included. As a complementary approach, we used CellChat, which infers ligand-receptor pairs and associated pathways based on a manually-curated list of literature-supported interactions grouped into 229 signaling pathways ^42^. Both approaches were consistent and showed that acinar and duct cells had the lowest number of enriched ligand and receptor genes compared to all other cell types while MECs had the highest number across epithelial populations (Figure S3A-C).

Differential expression analysis identified 9 ligand and 5 receptor genes among the *Etv1*+ cell-defining genes, as well as 5 ligands and 2 receptors in acinar cells (Figure 3A). The identified receptor genes enriched in *Etv1*+ cells included *Ghr*, *Dddr1*, *St14*, *Erbb3,* and *Epha5*, which were highly specific to this population (Figure 3B, left panel). On the other hand, the ligands found in *Etv1*+ cells were also enriched in other cell types, with the exception of *Col7a1,* which was highly specific (Figure 3B, right panel). A distinct set of ligands and receptors were enriched in acinar cells, including the receptor genes *Ntrk2* and *Kcnn4*, and the ligands *P4hb, Nucb2, Agt, Tcn2,* and *Pip.* All of the resulting putative interactions from our differential gene expression analysis are shown in Supplementary File 2. Interactions between acinar and *Etv1*+ cells with all other cell types are summarized as chord plots in Supplementary Figure S3D-E. All interactions predicted by CellChat are available in Supplementary File 3. Based simply on the total number of possible pairs (without accounting for the level of expression of individual genes), the strongest outgoing interactions from *Etv1*+ cell ligands were predicted to occur with receptors in endothelial cells, whereas *Etv1*+ cell receptors corresponded to ligands from myoepithelial and stromal cells. In contrast, the corresponding pairs for acinar cell ligands were expressed primarily in T-cells (Supplementary Figure S3D). These findings were largely corroborated by CellChat (Figue 3D), which does take into account the level of gene expression as well as the proportion of cells expressing a given ligand-receptor pair in a cluster.

**Figure 3.**
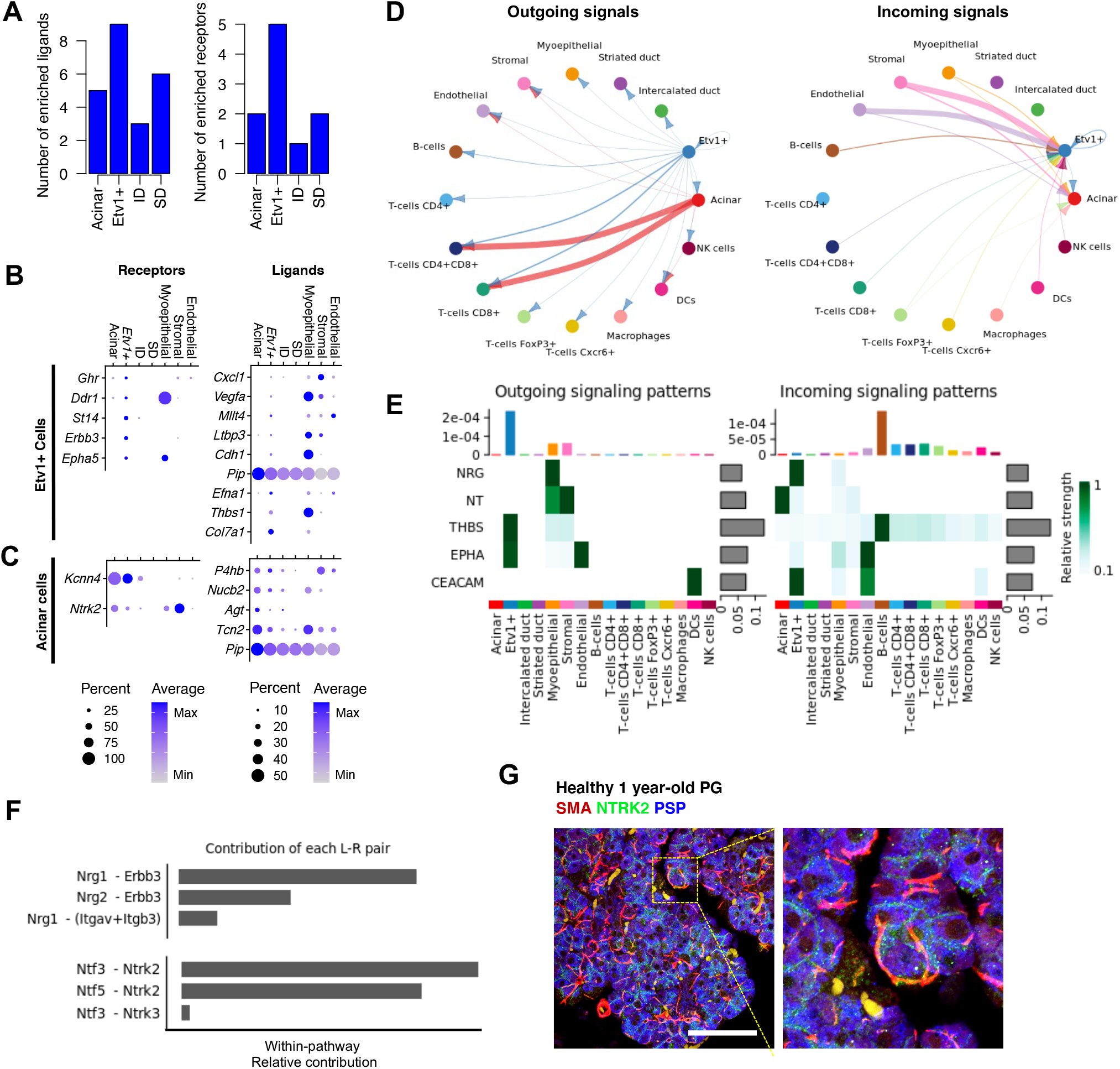
Ligand-receptor analysis of Etv1+ and acinar cells. A) Bar graphs with number of identified ligands and receptors among cell-defining genes from epithelial populations. B) Balloon plots of expression of ligands and receptors enriched in Etv1+ cells. C) Balloon plots of expression of ligands and receptors enriched in acinar cells. D) Chord plots summarizing putative ligand-receptor interactions with Acinar and Etv1+ cells, as predicted by CellChat. The arrows point to the cell expressing the receptors and are colored based on the source of the ligand. The thickness of the arrow is relative to the number of putative pairs identified. E) Heatmap shows the relative importance of each cell group based on CellChat-computed network centrality measures of NRG, NT, THBS, EPHA, and CEACAM signaling networks. F) Relative contribution of each ligand-receptor pair to the overall communication network of NRG and NT signaling pathways, which is the ratio of the total communication probability of the inferred network of each ligand-receptor pair to that of the signaling pathway. G) Immunofluorescence staining for Smooth muscle actin (SMA, Red), NTRK2 (green) and Parotid Secretory Protein (PSP, blue). The area delineated by the yellow dotted line is magnified to the right for visualization. Scale bar = 50um.

CellChat analysis determined that Etv1+ cells were a source of ligands for Thrombospondin (THBS) and EphrinA (EPHA) signaling pathways, and had receptors for Carcinoembryonic antigen cell adhesion molecule (CEACAM) and Neuregulin (NRG) ligands, whereas acinar cells were receptive to Neurotrophin (NT) signaling (Figure 3E). These results revealed notable interactions between myoepithelial and Etv1+ cells via the *Erbb3* receptor and two of its ligands, Neuregulin1 (*Nrg1*) and *Nrg2*, and between myoepithelial and acinar cells via the neurotrophin receptor *Ntrk2* and its ligands, Neurotrophin 3 (*Ntf3*) and *5* (*Ntf5*) (Figure 3E-F, Supplementary Figure S3D-E). *Ntrk2* was also expressed in *Etv1*+, myoepithelial and stromal cells in our scRNAseq data but immunofluorescence staining confirmed enrichment of the receptor in acinar cells of mouse parotid gland (Figure 3F). The cellular functions of *Ntrk2* in acinar cells are currently unknown and thus further mechanistic studies are warranted.

### CD8+CD4+ T-cells and acinar cells have the greatest transcriptional response to IR

The model of SG IR used in this study is based on a fractionated dosing schedule of 6Gy x 5 consecutive days, which leads to significant loss of saliva ^35^ but it does not result in extensive loss of acinar cells and development of fibrosis (Figure 4A) as reported by Ferreira *et al.* While this model shows a milder phenotype compared to alternative models using a single 15Gy dose in distinct mouse strains ^10, 43–45^, it has been previously used to demonstrate the therapeutic potential of adenovirus-based Neurturin gene transfer in the SMG to prevent the loss of saliva caused by IR. Given that we did not perform multiple technical replicates of each treatment, potential changes in cell proportions are reported as trends. In general, B cells and T cells were the most affected (Figure 4A-B). We observed a 33 % relative decrease in the proportion of B cells, a 39 % increase in CD4+ T cells, and a 195% increase in CD4+CD8+ T cells. A 22 % decrease in the proportion of acinar cells was also noted.

**Figure 4.**
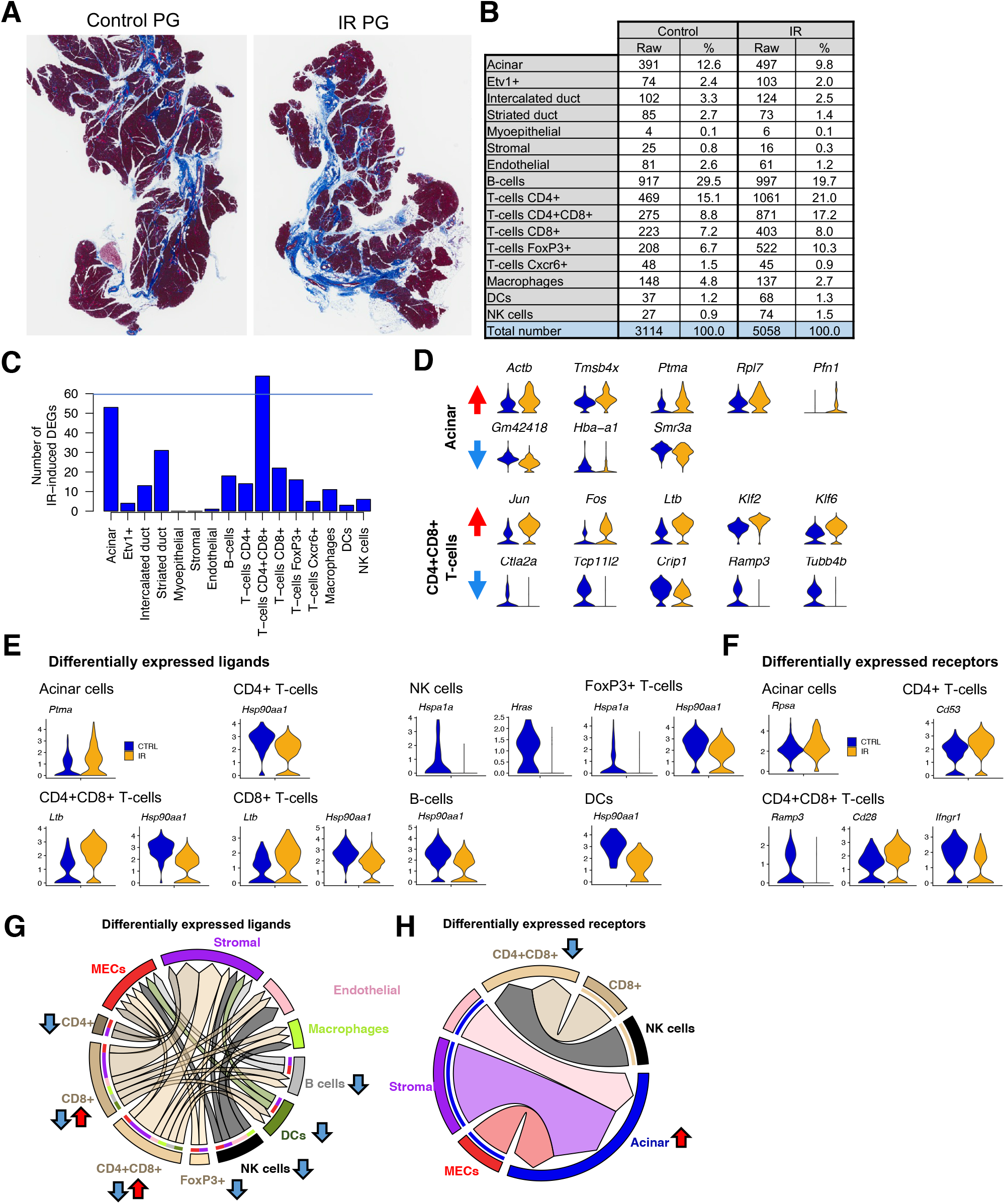
Cell-specific IR-induced DEGs. A) Masson’s trichrome staining of control and IR-PG collected 10-months post-IR. Areas in blue represent fibrotic/collagenous tissue. B) Cell numbers and proportions in scRNAseq datasets from control and irradiated PG. C) Bar graph showing number of DEGs post-IR in individual cell populations. DE analysis was performed with SEURAT’s default Wilcoxon test (p<0.05). D) Violin plots of top 5 (if present) up and downregulated genes in acinar and CD4+CD8+ T-cells. Red and blue arrows denote upregulated and downregulated genes, respectively. E-F) Violin plots of differentially expressed ligands and receptors from SEURAT’s DE analysis. G-H) Chord plots of ligand-receptor interactions with IR-induced DE receptors and ligands.

Differential expression analysis with SEURAT was performed between control and irradiated cell types. The complete list of differentially expressed genes (DEGs) is shown in Supplementary file 4. CD4+CD8+ T-cells had the highest number of dysregulated genes (∼70) post-IR across all identified cell populations, followed by acinar cells (Figure 4C). We did not detect DEGs in MEC and stromal populations post-IR, and only 1 gene was differentially expressed in IR endothelial cells. The lack of DEGs in MECs is likely explained because of the low number of MECs analyzed (Figure 4B). Stromal and endothelial cells also did not show significant changes in gene expression, but they were well-represented in our dataset; thus, cell numbers alone are not likely to account for the lack of DEGs post-IR in these populations. Instead, the lack of DEGs may reflect the fact that this model of IR damage is not highly fibrotic (Figure 4A). Alternatively, it’s possible that endothelial and stromal populations may have recovered a year after IR damage.

The top upregulated genes in acinar cells post-IR included *Actb, Tmsb4x*, and *Pfn1* which are involved in actin polymerization (Figure 4D). The genes *Gm42418, Hba-a1*, and *Smr3a* were the only downregulated genes in acinar cells and they were also downregulated in most other cell types (Figure S4A, Supplementary file 4), suggesting a global response to IR rather than an acinar-specific one. In CD4+CD8+ T-cells, the top upregulated genes post-IR were *Jun, Fos, Ltb*, *Klf2*, and *Klf6*, and the most downregulated genes were *Ctla2a, Tcp11l2, Crip1, Ramp3,* and *Tubb4b* (Figure 4D). In general, DEGs in acinar cells were associated with regulation of transepithelial transport, electron transport, apoptosis, and translation processes according to gene ontology analysis via The Gene Ontology Consortium ^46^, while DEGs in CD4+CD8+ T-cells were associated with V(D)J recombination, lymphocyte differentiation, apoptosis, axonogenesis, and ERK signaling pathway (Figure S4B).

When we cross-referenced the IR DEGs against the database of ligand-receptor pairs from Ramilowsky *et al.*, only a handful of ligands and receptors were represented (Figure 4E-F), and only a few of these had a corresponding pair (Figure 4G-H, Supplementary File 5). Most differentially expressed pairs were found between immune populations, MECs, stromal, and endothelial cells. In acinar and CD4+CD8+ T-cells, which were the most transcriptionally affected, we identified 5 ligands (*Ptma, Hsp90aa1, Ltb, Hspa1a,* and *Hras*) and 5 receptor genes (*Rpsa*, *Cd53, Ramp3, Cd28*, and *Ifngr1*) differentially expressed post-IR (Figure 4E-F). However, these DEGs were expressed across multiple clusters and were not defining for any individual population. For instance, *Hsp90aa1* was downregulated in all immune populations except NK cells and macrophages, and both *Hspa1a* and *Hras* were downregulated in NK cells (Figure 4E). Similarly, *Rpsa* was upregulated in acinar cells while *Ifngr1* was downregulated in CD4+CD8+ T-cells post-IR (Figure 4F). Putative pairs were found for *Rpsa* (Ribosomal protein SA (*Rpsa*), also known as Laminin receptor 1)*, Ifngr1* (Interferon Gamma Receptor 1)*, Hsp90aa1* (Heatshock protein 90 Alpha Family Class A Member 1)*, Ltb* (Lymphotoxin Beta), and *Hras* oncogene (Supplementary File 5).

This analysis suggested multiple signaling alterations including interactions with acinar cells via *Lamb2-Rpsa* and between NK and CD8+ cells with CD4+CD8+ T-cells via *Ifng-Ifngr1* (Supplementary File 5, Figure 4G-H). Paracrine signaling via *Hsp90aa1* from immune cells to *Egfr* expressed in myoepithelial, stromal, and endothelial cells was potentially reduced, while *Ltb* interaction with *Tnfrsf1a* and *Cd40* expressed by macrophages, endothelial cells, dendritic cells, and B-cells was potentially increased.

### Neurotrophin, neuregulin, ECM, and immune signaling are the main altered pathways in Acinar and Etv1+ cells post-IR

Given that too few ligands and receptors were differentially expressed, we next used CellChat, which is sensitive to expression patterns in ligands and receptors themselves, as well as their cofactors, and weighs the size of a given cluster and the proportion of cells within a cluster expressing a gene. CellChat predicted similar ligand-receptor interactions in the IR-PG compared to those from the control glands (Supplementary File 3). There was an increase in the number of interactions post-IR from 3128 in the control to 3191 in IR PG, although the predicted interaction strength was slightly reduced (Figure 5A). Interactions with *Rpsa*, *Hsp90aa1*, or *Hras* could not be confirmed since these are not included in CellChat’s database of ligand-receptor pairs. CellChat predicted that MECs, stromal, and endothelial cells had the highest number of differential interactions while B-cells and T-cells had the greatest difference in interaction strengths (Figure 5B, Figure S5A). A summary of the number of differential interactions per cell type is provided in Supplementary File 6. Here, we focus specifically on acinar and Etv1+ cells.

**Figure 5.**
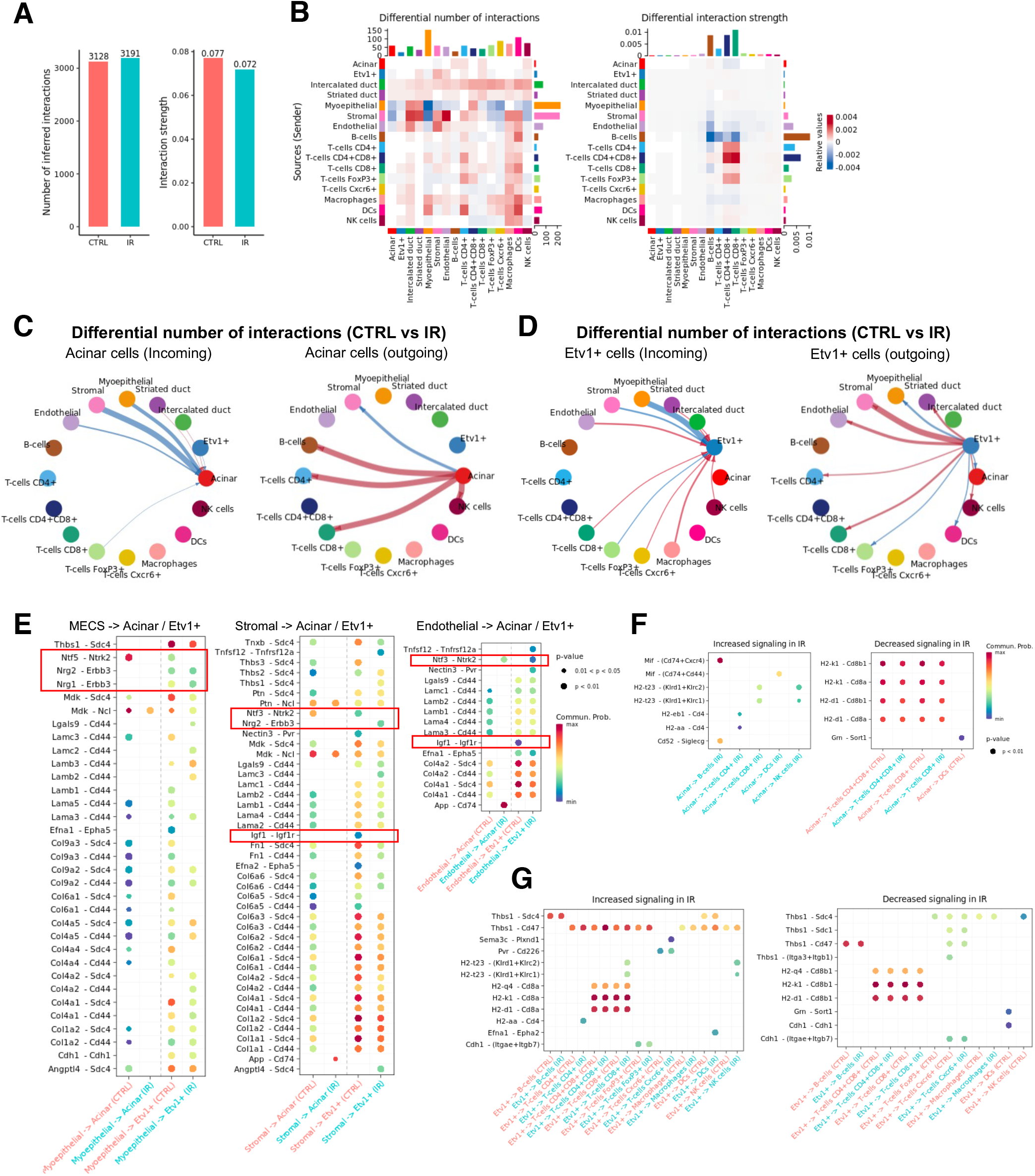
Dysregulated ligand-receptor pairs post-IR. A) Number of ligand-receptor pairs and interaction strength in control and IR-PG determined by CellChat. B) Differential number of possible interactions between acinar and Etv1+ cells with all other populations in IR-PG compared to control. Red (positive values) and blue (negative values) in the color bar indicate higher number of predicted interactions in IR-PG and controls, respectively. C-D) Differential ligand-receptor interactions in Acinar and Etv1+ cells between IR-PG and control, as predicted by CellChat. The arrows point to the cell expressing the receptors. Red arrows indicate increased number of interactions and blue arrows represent a decrease. The thickness of the arrow is relative to the change in number of interactions between control and IR-PG. E) Comparison of the significant ligand-receptor pairs between control and IR-PG, which contribute to the signaling from MECs, stromal and endothelial cells to acinar and Etv1+ cells. Dot color reflects communication probabilities and dot size represents computed p-values. Empty space means the communication probability is zero. p-values are computed from one-sided permutation test. F-G) Comparison of the significant ligand-receptor pairs between control and IR-PG, which contribute to the signaling from acinar and Etv1+ cells to immune populations. Dot color reflects communication probabilities and dot size represents computed p-values. Empty space means the communication probability is zero. p-values are computed from one-sided permutation test.

CellChat predicted altered signaling from MECs, stromal, and endothelial cells to both acinar and Etv1+ cells (Figure 5C-D). Generally, this seemed to be predominantly mediated by fewer interactions between *Cd44* and *Sdc4* with multiple collagens and laminins (Figure 5E). We also saw a shift in Neurotrophin signaling interactions post-IR, which were decreased between acinar cells and MECs or Stromal cells, but increased with endothelial cells (Figure 5E); this shift also involved changes in the ligand *Ntf5* and the appearance of a more significant contribution from *Ngf-Ntrk1* interactions (Figure S5B). Disruption of neurotrophin signaling was also recently reported in irradiated human salivary glands ^47^. Altered signaling in Etv1+ cells also involved loss of *Igf1-Igf1r* interactions with stromal and endothelial cells, and increased interactions via *Nrg2-Erbb3* with stromal cells (Figure 5E). A key difference between the two cell types was their differential interactions post-IR with immune populations. In acinar cells, there were increased interactions with T-cells and B-cells (Figure 5C, Figure 5F), but the strength of the interactions was generally lower (Figure S5C). In contrast, Etv1+ cells showed generally stronger interactions with T-cells, with the exception of FoxP3+ cells, which had fewer interactions (Figure 5D, Figure 5G, Figure S5C).

These results combined suggest that this model of IR injury shows alterations in cell adhesion interactions with the extracellular matrix (i.e. via Cd44, Sdc4, collagen and laminin), as well as changes in neurotrophin (Ntrk2, Ntf5, Ntf3), neuregulin (Nrg1, Nrg2, Erbb3), and IGF (Igf1, Igf1r) pathways. Furthermore, they suggest active involvement of immune cells, particularly T-cells, in mediating cellular responses post-IR, although the specific mechanisms involved are still unclear. Further mechanistic studies are encouraged to determine the functional relevance of these predicted interactions.

## Discussion

We generated a scRNAseq resource of adult PG that includes a chronic IR injury model. This resource allowed us to identify a discrete cell cluster of secretory cells defined by *Etv1* expression, and to predict putative ligand-receptor interactions that mediate key signaling pathways between secretory cell types and their microenvironment during homeostasis and post-injury.

The near exclusivity of *Etv1* expression in a single cluster is intriguing but it is not known whether it represents a cell-type-specific marker or a cell state. Their mixed histological localization suggests the latter. We saw expression of *Etv1* in both acinar and duct compartments, particularly close to the intercalated duct, similar to observations made by Song *et al.* in adult mouse SMG ^48^. *Etv1* is one of the top transcription factors in the salivary glands ^39^ and it was previously reported to be enriched in putative salivary stem cells defined by expression of Lin–CD24+c-Kit+Sca1+ ^49^. *Etv1* was also recently associated with the development of the acinar epithelium in the mouse SMG ^26^. These observations may be suggestive of *Etv1* being involved in an intermediate state between intercalated duct and acinar cells. Indeed, experiments in rodents suggest that intercalated duct cells may harbor stem cells that can differentiate into acinar cells or other duct cells ^50–52^. *Etv1+* cells showed enrichment of *Erbb3* expression, which was predicted to mediate signaling via neuregulin ligands derived from MECs and stromal cells. Erbb3 signaling is critical for SG development and plays a crucial role in organogenesis. Branching morphogenesis in the mouse SMG depends on intraepithelial signaling mediated by ErbB2, ErbB3, and neuregulin 1 (Nrg1) ^53^. Nrg1-null embryos show reduced innervation and defective branching morphogenesis ^54, 55^. Thus, it is plausible that *Etv1*+ (*Erbb3+)* cells in the adult parotid gland could be involved in either replenishment of the epithelium or wound healing, and may function as a proacinar population in the PG.

Our finding that the neurotrophic receptor *Ntrk2* is enriched in acinar cells is interesting because of the precedent of using neurotrophic factors such as neurturin to preserve function in irradiated SGs ^35, 56^. Moreover, we recently reported that in humans, IR chronically dysregulates the neurotrophin signaling pathway in both PG and SMG and is associated with functional and morphological abnormalities ^47^. Ligand-receptor analysis predicts that stromal cells and MECs communicate with *Ntrk2-*expressing acinar cells via *Ntf5* and *Ntf3*, respectively. Considering the localization of MECs surrounding acinar cells, it is likely that both juxtracrine and paracrine signaling takes place. The function of the Ntrk2 receptor in salivary acinar cells is not known but the gene is also highly expressed in Neurogenin 3-positive (Ngn3+) endocrine progenitors in the pancreas ^57^ and its activation regulates Ngn3+ cell fate commitment. Neurotrophin receptors are also mutated or upregulated in a variety of cancers, suggesting a role in proliferation and differentiation. In the SMG, *Ntrk2* is expressed in serous acinar cells but not in seromucous acinar cells ^26^, indicating that *Ntrk2* signaling may be important for the serous acinar phenotype, which is predominant in the PG. Furthermore, we recently identified that *NTRK2* is highly upregulated in MECs of irradiated human SGs along with other neurotrophin receptors and stimulation of neurotrophin signaling *in vitro* promoted myoepithelial differentiation^47^. In the lacrimal gland, neurotrophins are expressed in acini while neurotrophin receptors are expressed by MECs ^58^, suggesting that neurotrophin signaling may mediate intercellular communication between acinar cells and MECs in other exocrine tissues. Moreover, given that Ntrk2 is expressed on the cell surface, it may also provide a viable strategy to FACS-sort acinar cells from parotid gland to investigate expansion or differentiation of acinar cells *in vitro.* The latter application would likely require a combination of markers since *Ntrk2* is also expressed in *Etv1*+, myoepithelial and stromal cells.

The IR model used in our study has been previously used to demonstrate the protective effect of neurturin gene-transfer to prevent loss of saliva in the IR-SMG ^35^. In contrast to more severe models of IR injury ^10, 43, 45, 59, 60^, this model does not result in extensive loss of acinar cells and is only mildly fibrotic; thus, it is ideal for understanding transcriptional changes in acinar cells before they are lost. Indeed, acinar cells had the second largest number of DEGs post-IR in this model. Surprisingly, CD4+CD8+ cells showed the highest number of DEGs post-IR, suggesting that chronic damage post-IR may be sustained by immunologic mechanisms. There is growing evidence of immune-epithelial interactions in the regulation of tissue homeostasis and wound healing responses with macrophages and regulatory T-cells (T_regs_; FoxP3+) garnering the most attention ^61^. Through Notch-mediated signaling, mammary gland stem cells induced resident macrophages to produce Wnt ligands ultimately leading to mammary stem cell proliferation ^62^. Depletion of T_regs_ in the intestine leads to a reduction in LGR5+ stem cells ^63^. Given the extensive ligand-receptor interactions between *Etv1*+ cells and immune cells, it is interesting to speculate a functional role of *Etv1*+ cells in directing the localization and activation of resident immune populations. In the epidermis, distinct cellular populations around the hair follicle produce distinct chemokines to direct innate immune cell populations ^64^. For example, the interaction between *Etv1*+ and *FoxP3*+ cells via *Cdh1*-*Itgae* (E-cadherin and integrin-α-E) may represent the physical tethering of this sub-population of T-cells to the salivary epithelium under homeostasis ^65^, and it was increased post-IR (Figure 5G). It’s interesting to note that radiation treatment led to a 1.5-fold increase in T_regs_ without a concomitant change in *Etv1*+ cells or macrophages. Given the extensive role macrophages and *FoxP3*+ cells serve in injury and regeneration models, more work is required to unravel the impact of these T_regs_-epithelial interactions population during SG dysfunction.

Radiation treatment also resulted in the greatest increase in CD4+CD8+ populations and the most DEGs observed in the CD4+CD8+ cells (Figure 4). Clinical evaluation of SMG by immunohistochemistry following radiotherapy has revealed increased T-cells (CD3+, CD4+ or CD8+) in the periacinar area and B cell (CD20+) nodules in the periductal area ^66^. The DEGs in the CD4+CD8+ population suggest an imbalance in immune regulation following irradiation. Increases in KLF2 in IR PGs may represent a shift in T-cell populations as KLF2 is highly expressed in naïve and memory T-cells and downregulated by TCR activation and cytokine stimulation in effector T-cells ^67^. Additionally, high levels of KLF2 inhibit T-cell proliferation and clonal expansion ^67^. KLF6 also inhibits cell proliferation and is co-regulated with KLF2 in MCF-7 cells ^68^. Thus, high levels of KLF2 and KLF6 coupled with a lack of cytokines and chemokines on the DEGs suggest that the increase in CD4+CD8+ T-cells may represent a naïve population; however further kinetic analysis is required. This is also supported by a decrease in *Ctla2a,* which encodes for a cysteine protease that serves an immunosuppressive function in retinal pigment epithelium ^69, 70^ and promotes the conversion of CD4+ T cells to Treg cells via Transforming Growth Factor Beta (TGFβ) signaling ^71^. Lymphotoxin-β (LT-β), encoded by *Ltb*, is a TNF family member cytokine that has been predominantly studied in development and organization of lymphoid tissues ^72^. LT-β can mediate both regeneration and chronic tissue injury in epithelial organs via nuclear factor-κB (NF-κB) pathway ^73, 74^. Blocking the LT-β receptor suppresses immune responses by modulating trafficking mechanisms and disrupts the progression of T1DM in NOD mice ^72^. It is interesting to speculate whether the increased LT-β interactions with *Tnfrsf1a* or *CD40* prevent the clearance of immune populations or maintenance of naïve T cells. *Ltb* is induced following oxidative stress ^75^ and has been proposed to enable communication between lymphocytes and stromal cells ^73^, findings that are corroborated by this work predicting increased interactions with stromal and immune cell populations post-IR (Figure 5).

### Limitations of the study

A caveat of this study is the lack of isolation of basal ducts and peripheral nerve cells during PG dissociation, which were not represented. Similar limitations have been reported in other scRNAseq studies working with adult tissues, which could potentially be overcome using single nuclei RNAseq analysis. Furthermore, although multiple biological replicates were used, they were pooled together during dissociation prior to sequencing, thus, cell proportion changes should be considered with caution.

## Acknowledgments

The authors thank the support from Dr. Daniel Martin, Dr. Robert Morell, and Dr. Erich Boger from the Genomics and computational biology core (GCBC) at NIDCR for contributing to library preparation and sequencing. This work used the NIDCR Veterinary Resources Core (ZIC DE000740-05) and computational resources of the NIH HPC Biowulf cluster (http://hpc.nih.gov). The GCBC funds were from the NIDCD Division of Intramural Research/NIH (DC000086 to the GCBC). The study was supported by the Intramural Research Program of the National Institute of Dental and Craniofacial Research, NIH.

## Author Contributions

Conceptualization, writing and editing, A.M.C, B.R, K.H.L; Methodology, A.M.C., B.R., M.C.P., GCBC; Software, A.M.C, GCBC; Resources, M.P.H, K.H.L., A.M.C; Visualization, A.M.C., B.R., M.C.P; Data curation, project administration, and supervision, A.M.C.

## Declaration of Interests

The authors declare no competing interests.

## STAR METHODS

### RESOURCE AVAILABILITY

#### Lead contact

#### Materials Availability

This study did not generate new unique reagents.

#### Data and Code Availability

The single-cell RNAseq libraries were deposited in GEO under accession number GSE223516. The code used for analysis is available in github: https://github.com/chiblya/scRNAseq_PG. All package versions are reported in the gitgub repository. Ready-to-use Seurat objects are also available via figshare: 10.6084/m9.figshare.20406219

### EXPERIMENTAL MODEL AND SUBJECT DETAILS

#### C3H mice and irradiation (IR) treatment

C3H female mice were used for the study and were housed at the NIDCR Veterinary Resource Core in accordance with IACUC guidelines. At 6-10 weeks of age, mice received fractionated IR treatment at 5 × 6 Gy (6 Gy/day for 5 days). Only head and neck area was irradiated by placing each animal in a specially built Lucite jig so the animal could be immobilized without the use of anesthetics. IR treatment was delivered with by an X-Rad 320ix system. The mice were housed in a climate- and light-controlled environment, and allowed free access to food and water for 10 months post-IR before scRNA-seq analysis.

### METHODS

#### Single-cell Dissociation

Parotid glands from 2 female mice per treatment were dissociated in a 15ml gentleMACS C tube with 5ml of digestion enzyme using the human tumor dissociation kit (#130-095-929, Miltenyi Biotech, Auburn CA) in RPMI 1640 w/L-Glutamine (Cell applications, Inc, USA). Cell dissociation was performed in a Miltenyi gentleMACS Octo Dissociator using the manufacturer’s preset 37C_h_TDK_2 program. Following dissociation, 5ml of RPMI media were added to the dissociated cells and centrifuged at 1100 rpm for 10 min. Cells were resuspended in RPMI 1640 w/L-Glutamine with 5% PenStrep (Gibco, USA) and washed twice with RPMI. Cells were passed through 70 µm filters between centrifugation steps. Single-cell dissociation was confirmed by microscopic examination and cell concentration determined with a Cellometer (Nexcelom Biosciences). Cell concentration was adjusted to 5×10^5^ – 1×10^6^ cells/ml prior to analysis with a 10X genomics Next GEM Chromium controller.

#### Library prep and sequencing

Single-cell RNA-seq library preparation was performed at the NIDCR Genomics and Computational Biology Core using a Chromium Single Cell v3 method (10X Genomics) following the manufacturer’s protocol. Pooled single-cell RNA-seq libraries were sequenced on a NextSeq500 sequencer (Illumina). Cell Ranger Single-Cell Software Suite (10X Genomics) was used for demultiplexing, barcode assignment, and unique molecular identifier (UMI) quantification using the mm10 reference genome (Genome Reference Consortium Mouse Build 38) for read alignment.

#### Computational analysis

Cell Ranger files were imported to SEURAT v3 using R & R Studio software and processed for clustering following their default pipeline. As a quality control measure, cells with fewer than 200 genes were not included in subsequent analyses, and those with >5% of UMIs mapping to mitochondrial genes were defined as non-viable or apoptotic and were also excluded. These metrics were based on our previous scRNAseq analysis of murine SMG ^26^. Normalization and scaling were performed following SEURAT’s default pipeline. Data from control and irradiated glands were bioinformatically integrated prior to assigning cell annotations. ‘Clustree’ package was used to determine an optimal resolution for clustering and the resulting clusters were annotated based on the expression of known cell type markers. Cell-defining genes were determined using the ‘FindAllMarkers’ function which uses a Wilcoxon Rank Sum statistical test for analysis. Only genes with adjusted p-values <0.05 were considered as cell-defining genes. To identify DEGs between treatments, each population was compared individually using the ‘FindMarkers’ function from SEURAT package.

##### Ligand-receptor analysis

A database of curated ligand-receptor pairs was downloaded from Ramikowski *et al.* (2015). We used scripted code in R to automate the search for ligand and receptor genes within our dataset and leverage that information against the curated database. Additionally, we used CellChat v.1.6.1 (Jin, S. *et al.,* 2021) to predict significant interactions and their associated pathways. Plots were generated using the ‘circlize’ and complexHeatmap packages in R. The code is available in https://github.com/chiblya/scRNAseq_PG.

###### Immunohistochemistry

PGs were fixed in 4% paraformaldehyde overnight at 4°C and dehydrated with 70% Ethanol prior to paraffin embedding. 5µm sections were deparaffinized with xylene substitute for 10 minutes and rehydrated with reverse ethanol gradient for 5 minutes each. Heat induced antigen retrieval was performed using a microwave maintaining sub-boiling temperature for 10 minutes in a pH 6.0 Citrate Buffer (#21545, EDM Millipore, Darmstadt, Germany) . Sections were washed for 5 minutes with 0.1% Tween20 (Quality Biological, Inc) in PBS 1X (PBST). M.O.M.® (Mouse on Mouse) Immunodetection Kit (Vector Laboratories, Burlingame, CA) was used to block non-specific sites for 1 hour at room temperature followed by overnight incubation with primary antibodies at 4°C. Tissue sections were washed 3 times for 5 minutes each with PBST and incubated in secondary antibodies and nuclear stain (Hoechst (Thermo Fisher Scientific, Marietta, OH)) at room temperature for 1 hour. Coverslips were mounted with Fluoro-Gel (Electron Microscopy Sciences, Hatfield, PA), and imaging was performed with a Nikon A1R confocal system.

###### Stitch analysis

Etv1+ cell defining genes from control parotid sample (Supplementary File 1) were directly imported into STITCH (http://stitch.embl.de/). For reproducibility, analysis was performed selecting a minimum interaction score of 0.7 and limited to less than 10 interactions.

---

**Figure S1:**
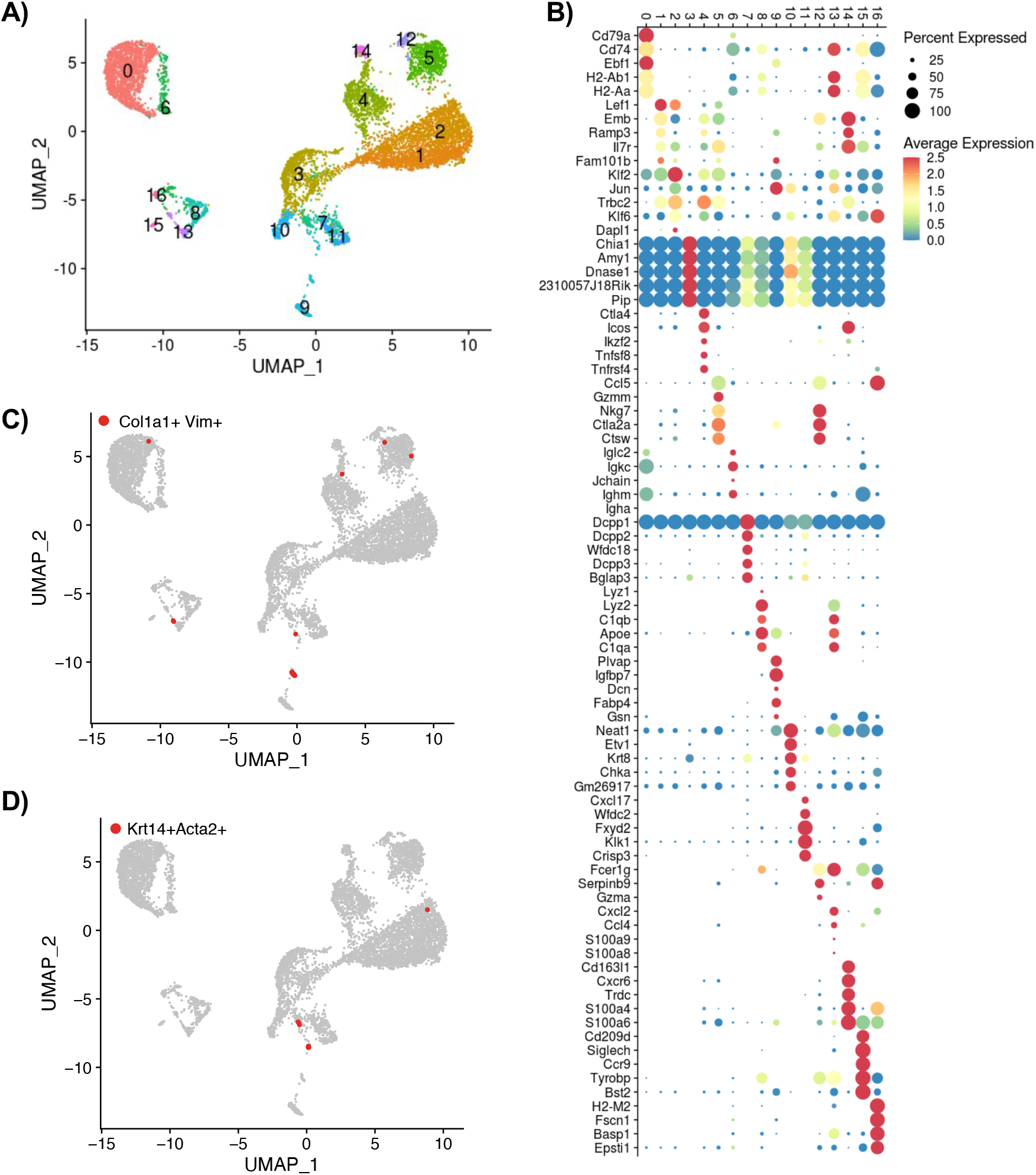
Annotation strategy. A) Unsupervised clustering of integrated control and irradiated mouse parotid gland (n=1 per treatment) B) Balloon plot of top cluster-defining genes. Color is relative to scaled gene expression and size of the dot represents the percentage of cells within a cluster expressing the gene C) UMAP highlighting cells that express the stromal markers Col1a1 and Vim D) UMAP highlighting myoepithelial cells that express Krt14 and Acta2

**Figure S2:**
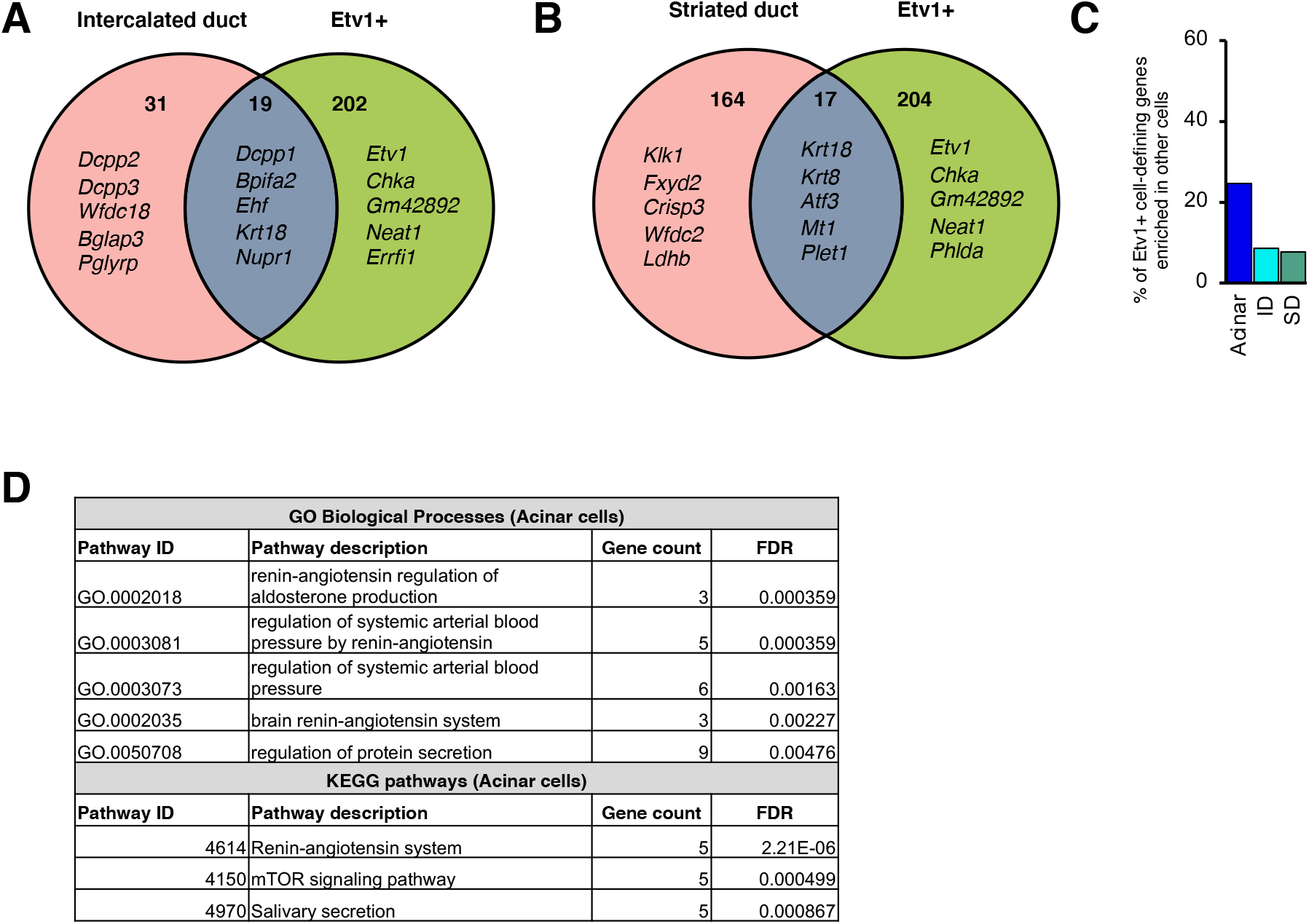
A) Venn diagram comparing defining genes for *Etv1*+ and ID populations. The numbers in the left and right panels indicates the number of unique genes in the corresponding population whereas the number in the central panel reflects the overlap between the two populations. B) Venn diagram comparing defining genes for *Etv1*+ and SD populations. The numbers in the left and right panels indicates the number of unique genes in the corresponding population whereas the number in the central panel reflects the overlap between the two populations. C) Bar graph with percentage of Etv1+ defining genes enriched in other epithelial cells. D) Results from STITCH analysis showing top biological processes and KEGG pathways associated with defining-genes from acinar cells.

**Figure S3.**
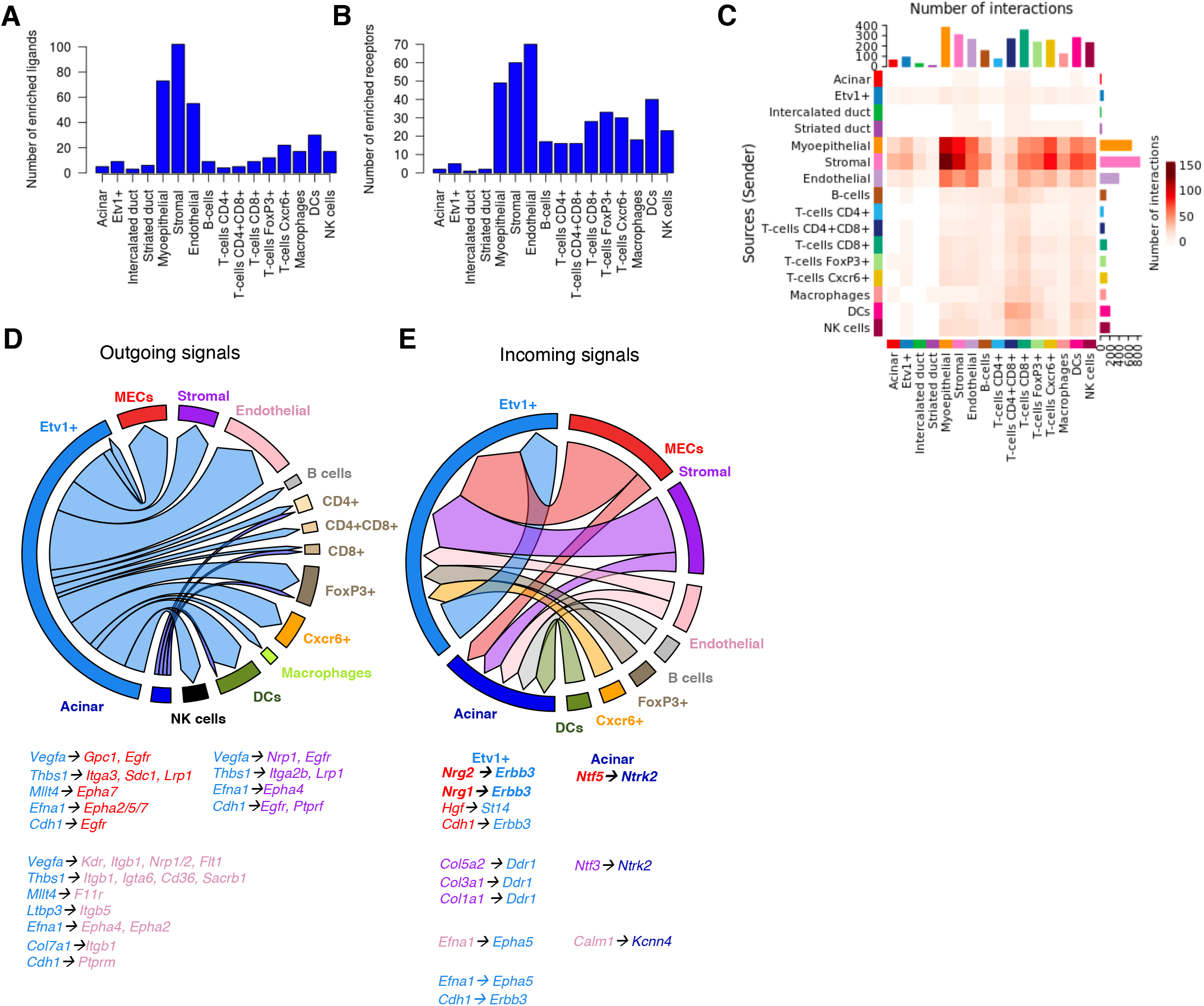
Ligand-receptor analysis of Etv1+ and acinar cells. A-B) Bar graphs with number of identified ligands and receptors among cell-defining genes from all populations. C) Heatmap of possible interactions between any two cell populations. Red (positive values) in the color bar indicate higher number of predicted interactions. D-E) Chord plot summarizing putative ligand-receptor interactions with Etv1+ cell ligands. The arrows point to the cell expressing the corresponding receptors and are color-coded based on the source of the ligand. The thickness of the arrow is relative to the number of putative pairs identified between Etv1 cells and the cell type pointed by the arrow. Representative ligand-receptor pairs are shown beside the chord plots.

**Figure S4.**
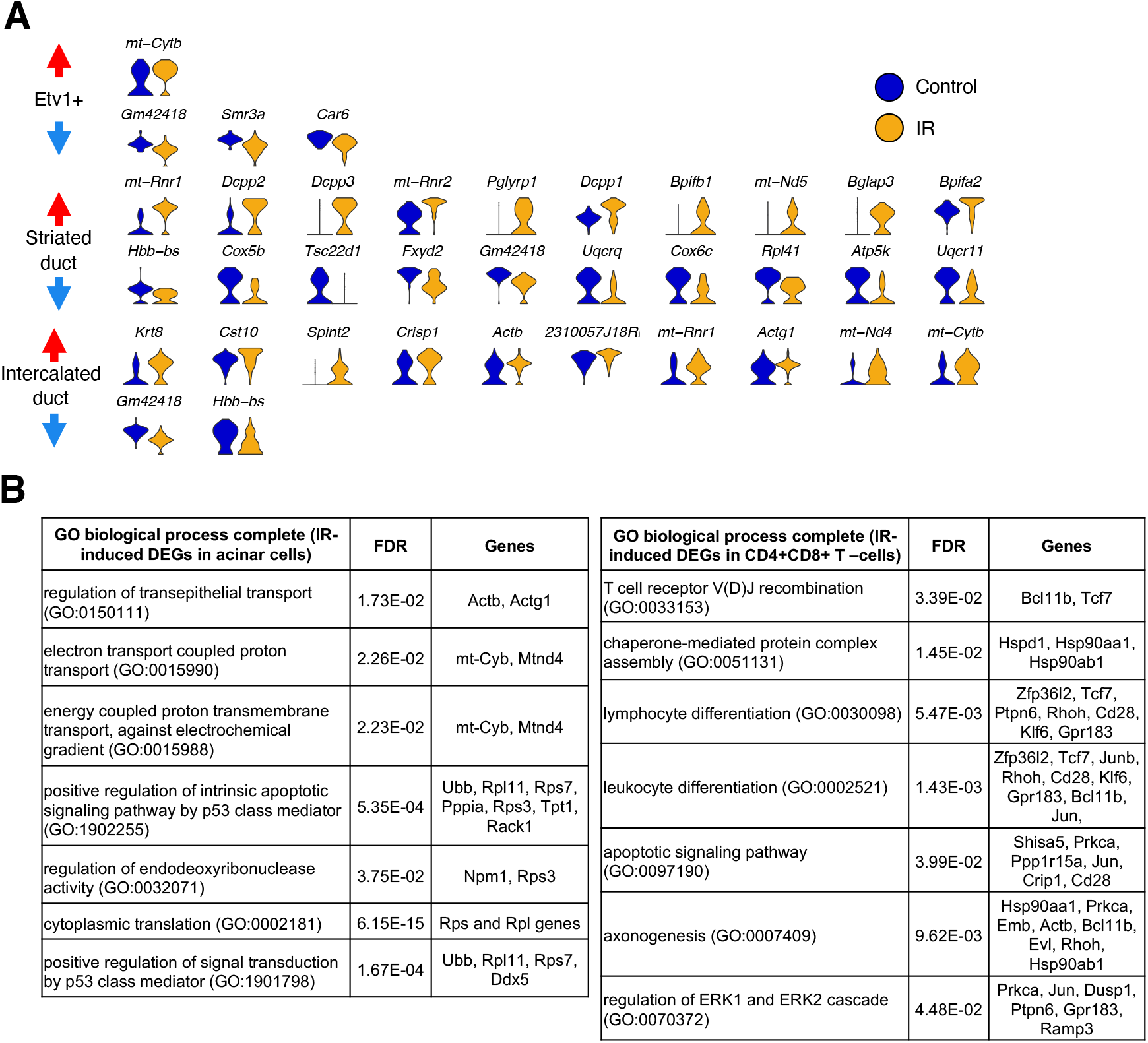
Cell-specific IR-induced DEGs. A) Violin plots of top 10 (if present) up and downregulated genes in epithelial populations. Red and blue arrows denote upregulated and downregulated genes, respectively. B) Representative output from gene ontology analysis with IR-induced DEGs in acinar and CD4+CD8+ T-cells showing dysregulated processes and their associated genes.

**Figure S5.**
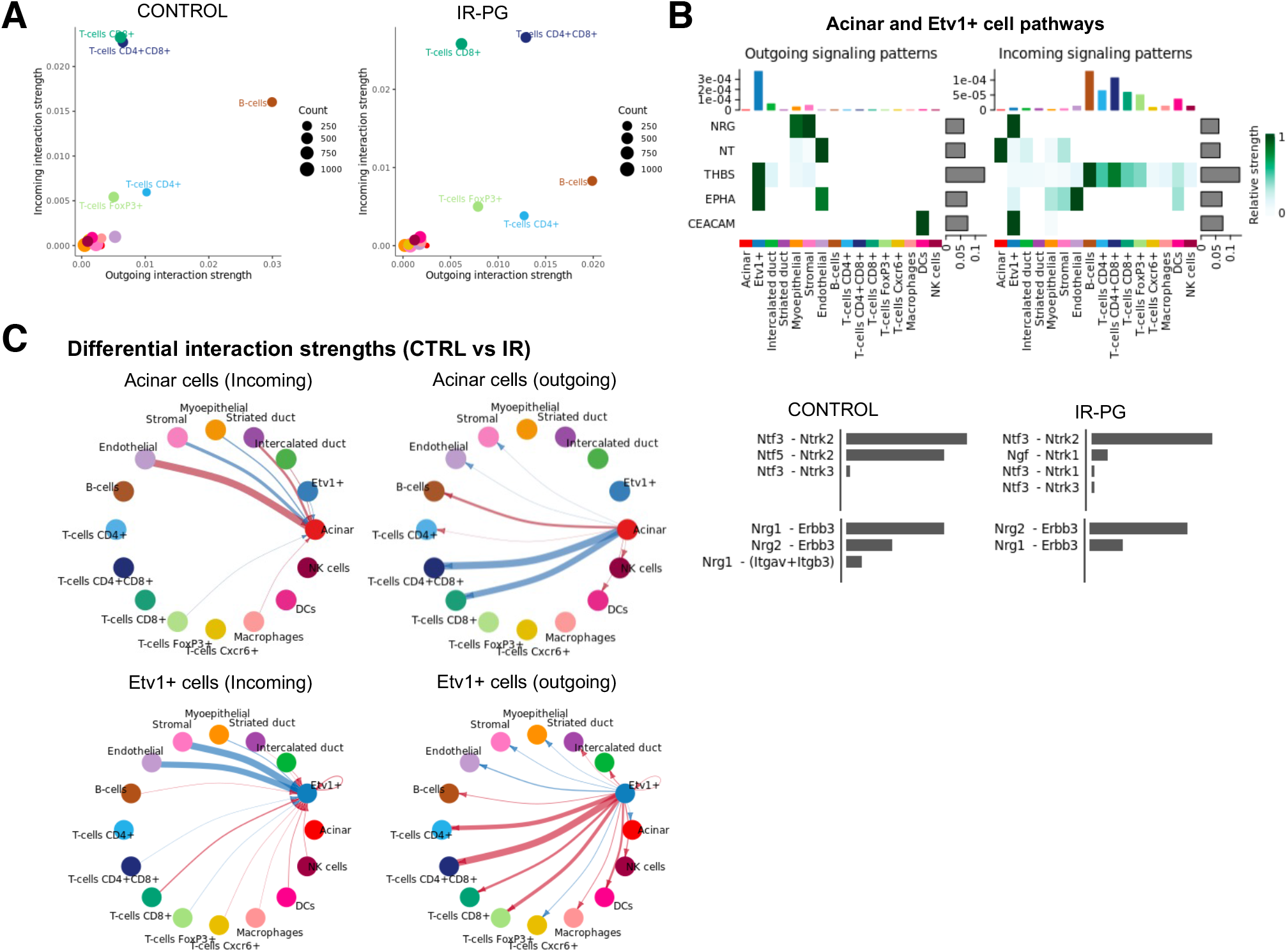
Dysregulated ligand-receptor pairs post-IR. A) 2D representation of incoming vs outgoing interaction strengths for all cell types relative to each other. B) Heatmap shows the relative importance of each cell group in IR-PG based on CellChat-computed network centrality measures of NRG, NT, THBS, EPHA, and CEACAM signaling networks. Relative contribution of each ligand-receptor pair to the overall communication network of NRG and NT signaling pathways, which is the ratio of the total communication probability of the inferred network of each ligand-receptor pair to that of the signaling pathway. C) Differential interactions strength between acinar and Etv1+ populations with all other cell types in IR-PG compared to control. Red (positive values) and blue (negative values) in the color bar indicate higher number of predicted interactions in IR-PG and controls, respectively

## Notes

### Competing Interest Statement

The authors have declared no competing interest.

### Summary of Updates

CellChat was used to complement the ligand-receptor analysis, and we have clarified the rationale and significance of the chosen irradiation model.

## References

1. Maruyama, C.L., Monroe, M.M., Hunt, J.P., Buchmann, L., and Baker, O.J. (2019). Comparing human and mouse salivary glands: A practice guide for salivary researchers. Oral Dis 25, 403–415. 10.1111/odi.12840.

2. Gao, X., Oei, M.S., Ovitt, C.E., Sincan, M., and Melvin, J.E. (2018). Transcriptional profiling reveals gland-specific differential expression in the three major salivary glands of the adult mouse. Physiological Genomics 50, 263–271. 10.1152/physiolgenomics.00124.2017.

3. Jensen, S.B., Pedersen, A.M., Vissink, A., Andersen, E., Brown, C.G., Davies, A.N., Dutilh, J., Fulton, J.S., Jankovic, L., Lopes, N.N., et al. (2010). A systematic review of salivary gland hypofunction and xerostomia induced by cancer therapies: prevalence, severity and impact on quality of life. Support Care Cancer 18, 1039–1060. 10.1007/s00520-010-0827-8.

4. Tasaka, S., Jingu, K., Takahashi, N., Umezawa, R., Yamamoto, T., Ishikawa, Y., Takeda, K., Suzuki, Y., and Kadoya, N. (2021). The Long-Term Recovery of Parotid Glands in Nasopharyngeal Carcinoma Treated by Intensity-Modulated Radiotherapy. Front Oncol 11, 665837. 10.3389/fonc.2021.665837.

5. Jensen, S.B., Vissink, A., Limesand, K.H., and Reyland, M.E. (2019). Salivary Gland Hypofunction and Xerostomia in Head and Neck Radiation Patients. J Natl Cancer Inst Monogr 2019. 10.1093/jncimonographs/lgz016.

6. Eisbruch, A., Dawson, L.A., Kim, H.M., Bradford, C.R., Terrell, J.E., Chepeha, D.B., Teknos, T.N., Anzai, Y., Marsh, L.H., Martel, M.K., et al. (1999). Conformal and intensity modulated irradiation of head and neck cancer: the potential for improved target irradiation, salivary gland function, and quality of life. Acta Otorhinolaryngol Belg 53, 271–275.

7. Henson, B.S., Eisbruch, A., D’Hondt, E., and Ship, J.A. (1999). Two-year longitudinal study of parotid salivary flow rates in head and neck cancer patients receiving unilateral neck parotid-sparing radiotherapy treatment. Oral Oncology 35, 234–241. https://doi.org/10.1016/S1368-8375(98)00104-3.

8. Avila, J.L., Grundmann, O., Burd, R., and Limesand, K.H. (2009). Radiation-induced salivary gland dysfunction results from p53-dependent apoptosis. Int J Radiat Oncol Biol Phys 73, 523–529. 10.1016/j.ijrobp.2008.09.036.

9. Meyer, S., Chibly, A.M., Burd, R., and Limesand, K.H. (2017). Insulin-Like Growth Factor-1-Mediated DNA Repair in Irradiated Salivary Glands Is Sirtuin-1 Dependent. J Dent Res 96, 225–232. 10.1177/0022034516677529.

10. Liu, X., Cotrim, A., Teos, L., Zheng, C., Swaim, W., Mitchell, J., Mori, Y., and Ambudkar, I. (2013). Loss of TRPM2 function protects against irradiation-induced salivary gland dysfunction. Nat Commun 4, 1515. 10.1038/ncomms2526.

11. Gilman, K.E., Camden, J.M., Klein, R.R., Zhang, Q., Weisman, G.A., and Limesand, K.H. (2019). P2X7 receptor deletion suppresses gamma-radiation-induced hyposalivation. Am J Physiol Regul Integr Comp Physiol 316, R687–R696. 10.1152/ajpregu.00192.2018.

12. Jasmer, K.J., Gilman, K.E., Munoz Forti, K., Weisman, G.A., and Limesand, K.H. (2020). Radiation-Induced Salivary Gland Dysfunction: Mechanisms, Therapeutics and Future Directions. J Clin Med 9. 10.3390/jcm9124095.

13. Grundmann, O., Mitchell, G.C., and Limesand, K.H. (2009). Sensitivity of Salivary Glands to Radiation: from Animal Models to Therapies. Journal of Dental Research 88, 894–903. 10.1177/0022034509343143.

14. Dirix, P., Nuyts, S., and Van den Bogaert, W. (2006). Radiation-induced xerostomia in patients with head and neck cancer: a literature review. Cancer 107, 2525–2534. 10.1002/cncr.22302.

15. Radfar, L., and Sirois, D.A. (2003). Structural and functional injury in minipig salivary glands following fractionated exposure to 70 Gy of ionizing radiation: an animal model for human radiation-induced salivary gland injury. Oral Surg Oral Med Oral Pathol Oral Radiol Endod 96, 267–274. 10.1016/s1079-2104(03)00369-x.

16. Li, Y., Taylor, J.M., Ten Haken, R.K., and Eisbruch, A. (2007). The impact of dose on parotid salivary recovery in head and neck cancer patients treated with radiation therapy. Int J Radiat Oncol Biol Phys 67, 660–669. 10.1016/j.ijrobp.2006.09.021.

17. Vissink, A., Mitchell, J.B., Baum, B.J., Limesand, K.H., Jensen, S.B., Fox, P.C., Elting, L.S., Langendijk, J.A., Coppes, R.P., and Reyland, M.E. (2010). Clinical management of salivary gland hypofunction and xerostomia in head-and-neck cancer patients: successes and barriers. Int J Radiat Oncol Biol Phys 78, 983–991. 10.1016/j.ijrobp.2010.06.052.

18. Grün, D., and van Oudenaarden, A. (2015). Design and Analysis of Single-Cell Sequencing Experiments. Cell 163, 799–810. https://doi.org/10.1016/j.cell.2015.10.039.

19. Kolodziejczyk, Aleksandra A., Kim, J.K., Svensson, V., Marioni, John C., and Teichmann, Sarah A. (2015). The Technology and Biology of Single-Cell RNA Sequencing. Molecular Cell 58, 610–620. https://doi.org/10.1016/j.molcel.2015.04.005.

20. Trapnell, C. (2015). Defining cell types and states with single-cell genomics. Genome Res 25, 1491–1498. 10.1101/gr.190595.115.

21. Wang, Y., and Navin, Nicholas E. (2015). Advances and Applications of Single-Cell Sequencing Technologies. Molecular Cell 58, 598–609. https://doi.org/10.1016/j.molcel.2015.05.005.

22. Tabula Muris, C., Overall, c., Logistical, c., Organ, c., processing, Library, p., sequencing, Computational data, a., Cell type, a., Writing, g., et al. (2018). Single-cell transcriptomics of 20 mouse organs creates a Tabula Muris. Nature 562, 367–372. 10.1038/s41586-018-0590-4.

23. Tabula Sapiens, C., Jones, R.C., Karkanias, J., Krasnow, M.A., Pisco, A.O., Quake, S.R., Salzman, J., Yosef, N., Bulthaup, B., Brown, P., et al. (2022). The Tabula Sapiens: A multiple-organ, single-cell transcriptomic atlas of humans. Science 376, eabl4896. 10.1126/science.abl4896.

24. Chen, M., Lin, W., Gan, J., Lu, W., Wang, M., Wang, X., Yi, J., and Zhao, Z. (2022). Transcriptomic Mapping of Human Parotid Gland at Single-Cell Resolution. J Dent Res 101, 972–982. 10.1177/00220345221076069.

25. Huang, N., Perez, P., Kato, T., Mikami, Y., Okuda, K., Gilmore, R.C., Conde, C.D., Gasmi, B., Stein, S., Beach, M., et al. (2021). SARS-CoV-2 infection of the oral cavity and saliva. Nat Med 27, 892–903. 10.1038/s41591-021-01296-8.

26. Hauser, B.R., Aure, M.H., Kelly, M.C., Hoffman, M.P., and Chibly, A.M. (2020). Generation of a Single-Cell RNAseq Atlas of Murine Salivary Gland Development. iScience 23, 101838. https://doi.org/10.1016/j.isci.2020.101838.

27. Oyelakin, A., Song, E.A.C., Min, S., Bard, J.E., Kann, J.V., Horeth, E., Smalley, K., Kramer, J.M., Sinha, S., and Romano, R.A. (2019). Transcriptomic and Single-Cell Analysis of the Murine Parotid Gland. J Dent Res 98, 1539–1547. 10.1177/0022034519882355.

28. Sekiguchi, R., Martin, D., Genomics, Computational Biology, C., and Yamada, K.M. (2020). Single-Cell RNA-seq Identifies Cell Diversity in Embryonic Salivary Glands. J Dent Res 99, 69–78. 10.1177/0022034519883888.

29. Praktiknjo, S.D., Obermayer, B., Zhu, Q., Fang, L., Liu, H., Quinn, H., Stoeckius, M., Kocks, C., Birchmeier, W., and Rajewsky, N. (2020). Tracing tumorigenesis in a solid tumor model at single-cell resolution. Nat Commun 11, 991. 10.1038/s41467-020-14777-0.

30. Horeth, E., Oyelakin, A., Song, E.C., Che, M., Bard, J., Min, S., Kiripolsky, J., Kramer, J.M., Sinha, S., and Romano, R.A. (2021). Transcriptomic and Single-Cell Analysis Reveals Regulatory Networks and Cellular Heterogeneity in Mouse Primary Sjogren’s Syndrome Salivary Glands. Front Immunol 12, 729040. 10.3389/fimmu.2021.729040.

31. Hong, X., Meng, S., Tang, D., Wang, T., Ding, L., Yu, H., Li, H., Liu, D., Dai, Y., and Yang, M. (2020). Single-Cell RNA Sequencing Reveals the Expansion of Cytotoxic CD4(+) T Lymphocytes and a Landscape of Immune Cells in Primary Sjogren’s Syndrome. Front Immunol 11, 594658. 10.3389/fimmu.2020.594658.

32. Xu, Y., Feng, S., Peng, Q., Zhu, W., Zu, Q., Yao, X., Zhang, Q., Cao, J., and Jiao, Y. (2021). Single-cell RNA sequencing reveals the cell landscape of a radiation-induced liver injury mouse model. Radiation Medicine and Protection 2, 181–183. 10.1016/j.radmp.2021.11.001.

33. Mukherjee, A., Epperly, M.W., Shields, D., Hou, W., Fisher, R., Hamade, D., Wang, H., Saiful Huq, M., Bao, R., Tabib, T., et al. (2021). Ionizing irradiation-induced Fgr in senescent cells mediates fibrosis. Cell Death Discov 7, 349. 10.1038/s41420-021-00741-4.

34. Paldor, M., Levkovitch-Siany, O., Eidelshtein, D., Adar, R., Enk, C.D., Marmary, Y., Elgavish, S., Nevo, Y., Benyamini, H., Plaschkes, I., et al. (2022). Single-cell transcriptomics reveals a senescence-associated IL-6/CCR6 axis driving radiodermatitis. EMBO Mol Med, e15653. 10.15252/emmm.202115653.

35. Ferreira, J.N.A., Zheng, C., Lombaert, I.M.A., Goldsmith, C.M., Cotrim, A.P., Symonds, J.M., Patel, V.N., and Hoffman, M.P. (2018). Neurturin Gene Therapy Protects Parasympathetic Function to Prevent Irradiation-Induced Murine Salivary Gland Hypofunction. Mol Ther Methods Clin Dev 9, 172–180. 10.1016/j.omtm.2018.02.008.

36. Stuart, T., Butler, A., Hoffman, P., Hafemeister, C., Papalexi, E., Mauck, W.M., Hao, Y., Stoeckius, M., Smibert, P., and Satija, R. (2019). Comprehensive Integration of Single-Cell Data. Cell 177, 1888–1902.e1821. https://doi.org/10.1016/j.cell.2019.05.031.

37. Hao, Y., Hao, S., Andersen-Nissen, E., Mauck, W.M., 3rd, Zheng, S., Butler, A., Lee, M.J., Wilk, A.J., Darby, C., Zager, M., et al. (2021). Integrated analysis of multimodal single-cell data. Cell 184, 3573–3587 e3529. 10.1016/j.cell.2021.04.048.

38. Zappia, L., and Oshlack, A. (2018). Clustering trees: a visualization for evaluating clusterings at multiple resolutions. GigaScience 7. 10.1093/gigascience/giy083.

39. Michael, D.G., Pranzatelli, T.J.F., Warner, B.M., Yin, H., and Chiorini, J.A. (2019). Integrated Epigenetic Mapping of Human and Mouse Salivary Gene Regulation. J Dent Res 98, 209–217. 10.1177/0022034518806518.

40. Kuhn, M., von Mering, C., Campillos, M., Jensen, L.J., and Bork, P. (2008). STITCH: interaction networks of chemicals and proteins. Nucleic Acids Res 36, D684–688. 10.1093/nar/gkm795.

41. Ramilowski, J.A., Goldberg, T., Harshbarger, J., Kloppmann, E., Lizio, M., Satagopam, V.P., Itoh, M., Kawaji, H., Carninci, P., Rost, B., and Forrest, A.R. (2015). A draft network of ligand-receptor-mediated multicellular signalling in human. Nat Commun 6, 7866. 10.1038/ncomms8866.

42. Jin, S., Guerrero-Juarez, C.F., Zhang, L., Chang, I., Ramos, R., Kuan, C.H., Myung, P., Plikus, M.V., and Nie, Q. (2021). Inference and analysis of cell-cell communication using CellChat. Nat Commun 12, 1088. 10.1038/s41467-021-21246-9.

43. Teos, L.Y., Zheng, C.Y., Liu, X., Swaim, W.D., Goldsmith, C.M., Cotrim, A.P., Baum, B.J., and Ambudkar, I.S. (2016). Adenovirus-mediated hAQP1 expression in irradiated mouse salivary glands causes recovery of saliva secretion by enhancing acinar cell volume decrease. Gene Therapy 23, 572–579. 10.1038/gt.2016.29.

44. Zheng, C., Cotrim, A.P., Rowzee, A., Swaim, W., Sowers, A., Mitchell, J.B., and Baum, B.J. (2011). Prevention of radiation-induced salivary hypofunction following hKGF gene delivery to murine submandibular glands. Clin Cancer Res 17, 2842–2851. 10.1158/1078-0432.CCR-10-2982.

45. Lombaert, I.M., Brunsting, J.F., Wierenga, P.K., Kampinga, H.H., de Haan, G., and Coppes, R.P. (2008). Keratinocyte growth factor prevents radiation damage to salivary glands by expansion of the stem/progenitor pool. Stem Cells 26, 2595–2601. 10.1634/stemcells.2007-1034.

46. The Gene Ontology Consortium (2019). The Gene Ontology Resource: 20 years and still GOing strong. Nucleic Acids Research 47, D330–D338. 10.1093/nar/gky1055.

47. Chibly, A.M., Patel V. N., Aure M. H., Pasquale M. C., NIDCD/NIDCR Genomics and Computational Biology Core, Martin G. E., Ghannam M., Andrade J., Denegre N., Simpson C., Goldstein D. P., Liu F., Lombaert I. M. A., Hoffman, M. P. (2023). Neurotrophin signaling is a central mechanism of salivary dysfunction after irradiation that disrupts myoepithelial cells. NPJ Regenerative Medicine (Accepted, In press).

48. Song, E.A.C., Smalley, K., Oyelakin, A., Horeth, E., Che, M., Wrynn, T., Osinski, J., Romano, R.A., and Sinha, S. (2022). Genetic Study of Elf5 and Ehf in the Mouse Salivary Gland. J Dent Res, 220345221130258. 10.1177/00220345221130258.

49. Xiao, N., Lin, Y., Cao, H., Sirjani, D., Giaccia, A.J., Koong, A.C., Kong, C.S., Diehn, M., and Le, Q.T. (2014). Neurotrophic factor GDNF promotes survival of salivary stem cells. J Clin Invest 124, 3364–3377. 10.1172/JCI74096.

50. Katsumata, O., Sato, Y., Sakai, Y., and Yamashina, S. (2009). Intercalated duct cells in the rat parotid gland may behave as tissue stem cells. Anat Sci Int 84, 148–154. 10.1007/s12565-009-0019-0.

51. Denny, P.C., Liu, P., and Denny, P.A. (1999). Evidence of a phenotypically determined ductal cell lineage in mouse salivary glands. The Anatomical Record 256, 84–90. 10.1002/(sici)1097-0185(19990901)256:1&<84::Aid-ar11>3.0.Co;2-s.

52. Denny, P.C., Ball, W.D., and Redman, R.S. (1997). Salivary glands: a paradigm for diversity of gland development. Crit Rev Oral Biol Med 8, 51–75. 10.1177/10454411970080010301.

53. Miyazaki, Y., Nakanishi, Y., and Hieda, Y. (2004). Tissue interaction mediated by neuregulin-1 and ErbB receptors regulates epithelial morphogenesis of mouse embryonic submandibular gland. Dev Dyn 230, 591–596. 10.1002/dvdy.20078.

54. Nedvetsky, Pavel I., Emmerson, E., Finley, Jennifer K., Ettinger, A., Cruz-Pacheco, N., Prochazka, J., Haddox, Candace L., Northrup, E., Hodges, C., Mostov, Keith E., et al. (2014). Parasympathetic Innervation Regulates Tubulogenesis in the Developing Salivary Gland. Developmental Cell 30, 449–462. https://doi.org/10.1016/j.devcel.2014.06.012.

55. Mattingly, A., Finley, J.K., and Knox, S.M. (2015). Salivary gland development and disease. Wiley Interdiscip Rev Dev Biol 4, 573–590. 10.1002/wdev.194.

56. Lombaert, I.M.A., Patel, V.N., Jones, C.E., Villier, D.C., Canada, A.E., Moore, M.R., Berenstein, E., Zheng, C., Goldsmith, C.M., Chorini, J.A., et al. (2020). CERE-120 Prevents Irradiation-Induced Hypofunction and Restores Immune Homeostasis in Porcine Salivary Glands. Molecular Therapy – Methods & Clinical Development 18, 839–855. https://doi.org/10.1016/j.omtm.2020.07.016.

57. Shamblott, M.J., O’Driscoll, M.L., Gomez, D.L., and McGuire, D.L. (2016). Neurogenin 3 is regulated by neurotrophic tyrosine kinase receptor type 2 (TRKB) signaling in the adult human exocrine pancreas. Cell Communication and Signaling 14, 23. 10.1186/s12964-016-0146-x.

58. Ghinelli, E., Johansson, J., R, J.D., Chen, L.-L., Zoukhri, D., Hodges, R.R., and Dartt, D.A. (2003). Presence and Localization of Neurotrophins and Neurotrophin Receptors in Rat Lacrimal Gland. Investigative Ophthalmology & Visual Science 44, 3352–3357. 10.1167/iovs.03-0037.

59. Lombaert, I.M., Brunsting, J.F., Wierenga, P.K., Faber, H., Stokman, M.A., Kok, T., Visser, W.H., Kampinga, H.H., de Haan, G., and Coppes, R.P. (2008). Rescue of salivary gland function after stem cell transplantation in irradiated glands. PLoS One 3, e2063. 10.1371/journal.pone.0002063.

60. Ninche, N., Kwak, M., and Ghazizadeh, S. (2020). Diverse epithelial cell populations contribute to the regeneration of secretory units in injured salivary glands. Development 147. 10.1242/dev.192807.

61. Naik, S., Larsen, S.B., Cowley, C.J., and Fuchs, E. (2018). Two to Tango: Dialog between Immunity and Stem Cells in Health and Disease. Cell 175, 908–920. https://doi.org/10.1016/j.cell.2018.08.071.

62. Chakrabarti, R., Celià-Terrassa, T., Kumar, S., Hang, X., Wei, Y., Choudhury, A., Hwang, J., Peng, J., Nixon, B., Grady, J.J., et al. (2018). Notch ligand Dll1 mediates cross-talk between mammary stem cells and the macrophageal niche. Science 360, eaan4153. 10.1126/science.aan4153.

63. Biton, M., Haber, A.L., Rogel, N., Burgin, G., Beyaz, S., Schnell, A., Ashenberg, O., Su, C.-W., Smillie, C., Shekhar, K., et al. (2018). T Helper Cell Cytokines Modulate Intestinal Stem Cell Renewal and Differentiation. Cell 175, 1307–1320. e1322. https://doi.org/10.1016/j.cell.2018.10.008.

64. Mansfield, K., and Naik, S. (2020). Unraveling Immune-Epithelial Interactions in Skin Homeostasis and Injury. Yale J Biol Med 93, 133–143.

65. Agace, W.W., Higgins, J.M., Sadasivan, B., Brenner, M.B., and Parker, C.M. (2000). T-lymphocyte-epithelial-cell interactions: integrin alpha(E)(CD103)beta(7), LEEP-CAM and chemokines. Curr Opin Cell Biol 12, 563–568. 10.1016/s0955-0674(00)00132-0.

66. Teymoortash, A., Simolka, N., Schrader, C., Tiemann, M., and Werner, J.A. (2005). Lymphocyte subsets in irradiation-induced sialadenitis of the submandibular gland. Histopathology 47, 493–500. 10.1111/j.1365-2559.2005.02256.x.

67. Preston, G.C., Feijoo-Carnero, C., Schurch, N., Cowling, V.H., and Cantrell, D.A. (2013). The Impact of KLF2 Modulation on the Transcriptional Program and Function of CD8 T Cells. PLOS ONE 8, e77537. 10.1371/journal.pone.0077537.

68. Ebert, R., Zeck, S., Meissner-Weigl, J., Klotz, B., Rachner, T.D., Benad, P., Klein-Hitpass, L., Rudert, M., Hofbauer, L.C., and Jakob, F. (2012). Krüppel-like factors KLF2 and 6 and Ki-67 are direct targets of zoledronic acid in MCF-7 cells. Bone 50, 723–732. https://doi.org/10.1016/j.bone.2011.11.025.

69. Sugita, S., Horie, S., Nakamura, O., Maruyama, K., Takase, H., Usui, Y., Takeuchi, M., Ishidoh, K., Koike, M., Uchiyama, Y., et al. (2009). Acquisition of T Regulatory Function in Cathepsin L-Inhibited T Cells by Eye-Derived CTLA-2α during Inflammatory Conditions. The Journal of Immunology 183, 5013. 10.4049/jimmunol.0901623.

70. Sugita, S., Horie, S., Nakamura, O., Futagami, Y., Takase, H., Keino, H., Aburatani, H., Katunuma, N., Ishidoh, K., Yamamoto, Y., and Mochizuki, M. (2008). Retinal Pigment Epithelium-Derived CTLA-2α Induces TGFβ-Producing T Regulatory Cells. The Journal of Immunology 181, 7525. 10.4049/jimmunol.181.11.7525.

71. Sugita, S., Yamada, Y., Horie, S., Nakamura, O., Ishidoh, K., Yamamoto, Y., Yamagami, S., and Mochizuki, M. (2011). Induction of T Regulatory Cells by Cytotoxic T-Lymphocyte Antigen-2α on Corneal Endothelial Cells. Investigative Ophthalmology & Visual Science 52, 2598–2605. 10.1167/iovs.10-6322.

72. McCarthy, D.D., Summers-Deluca, L., Vu, F., Chiu, S., Gao, Y., and Gommerman, J.L. (2006). The lymphotoxin pathway. Immunologic Research 35, 41–53. 10.1385/IR:35:1:41.

73. Wolf, M.J., Seleznik, G.M., Zeller, N., and Heikenwalder, M. (2010). The unexpected role of lymphotoxin β receptor signaling in carcinogenesis: from lymphoid tissue formation to liver and prostate cancer development. Oncogene 29, 5006–5018. 10.1038/onc.2010.260.

74. Tumanov, A.V., Koroleva, E.P., Christiansen, P.A., Khan, M.A., Ruddy, M.J., Burnette, B., Papa, S., Franzoso, G., Nedospasov, S.A., Fu, Y.X., and Anders, R.A. (2009). T Cell-Derived Lymphotoxin Regulates Liver Regeneration. Gastroenterology 136, 694–704.e694. https://doi.org/10.1053/j.gastro.2008.09.015.

75. Wong, G.H.W. (1995). Protective roles of cytokines against radiation: Induction of mitochondrial MnSOD. Biochimica et Biophysica Acta (BBA) – Molecular Basis of Disease 1271, 205–209. https://doi.org/10.1016/0925-4439(95)00029-4.

